# A cost-effective way to design a two layered cell trapping microfluidic device

**DOI:** 10.1101/2021.05.29.446315

**Authors:** José M. Lázaro-Guevara

## Abstract

When studying cells population, sometimes is necessary to isolate a specific subset to perform a timeline analysis or determinate the evolution on cell progeny. Microfluidic devices are being used for fulfill this function in multiple designs. However, the design and fabrication of this type of devices involve multiple challenges and high cost. Miniaturization tends to be the primarily challenge where those devices below 10um tends to be so expensive that needs special facilities for fabrication, but using techniques as multilayer fabrication those problems can be overcome. In this work are stated techniques (photolithography, soft-lithography) that helps solving that limitations, from using inexpensive materials, to alignment techniques that does not requires high-tech equipment’s, and a practical design for microfluidic cell trapping array. Where multiple cell types could be seeded and trapped in small cluster (varying on cell size) to be retrieved later on in an easy and practical way.

## 1. Introduction

When studying cells population, sometimes is necessary to isolate a specific subset to perform a timeline analysis or determinate the evolution on cell progeny. Microfluidic devices are being used for fulfill this function in multiple designs. (Kimmerling et al., 2016; Lin, Chu, Thiery, Lim, & Rodriguez, 2013)

However, those devices tend to be designed using high-resolution photolithographic techniques that implies extreme accuracy and expensive materials.(Qin, Xia, & Whitesides, 2010)

Obtaining the same results, using inexpensive material is a complicated task, based on the limitation on printing possibilities under 10 microns using non-chrome photomask at 25,000 dpi, and it is at this tiny size where these cell-trap devices fulfill their function.

To overcome the limitations a new approach has been used, whereas the width limitation using a regular photomask is 10 microns, and the thickness limitation is around 2 microns; allowing a multilayer design that compensates the lack of miniaturization on width.(Choi & Park, 2010) In the design of a cell-trapping device, a functional system could be achieved, using multi-step lithography that consist in overpassing a silicon wafer over the lithographic phase as many times as needed, but for a cell trap array only two layers of 5 and 15 microns are sufficient to achieve this goal. (Alvaro, Aaron, & Shuvo, 2006)

## 2. Materials and Methods

### 2.1 Photolithography

Photolithography, optical lithography or UV-lithography, is a process used in microfabrication to pattern thin film designs over a substrate, using light to transfer the pattern from a photomask to a material covered with light-sensitive resin. (Jaeger, 2002)

#### 2.1.1 Design and photomask

The early stages of the design were focused to determinate the form and how many layers were needed for this specific device, we used as a reference model the one proposed by Kimmerling et al. (Fig 1 A). However, based on the need of 3 microns channels was impossible to replicate with the equipment in our laboratory, so we created a multilayer design (Fig 1 B). The drawing was transcribed from paper version to 2D-CAD model using DraftSight software (Fig 1 C). Then the design undergone multiple changes to improve the possibilities of succeeding in functionality, and spliced into two different-layers (Fig 1 D; E), the variations in design cover from tolerance between layers ranging from 50 to 300 micrometers into connecting channels (Fig 1 E), as size of the trapping cell chamber that also ranges from 50 to 300 micrometers. Finally the 2D draw was sent to outputcity.com for get printed at 25,000 dpi for a negative photoresist resin (Fig 1 F), finally obtaining two photomask for next process.

**Figure 1.**
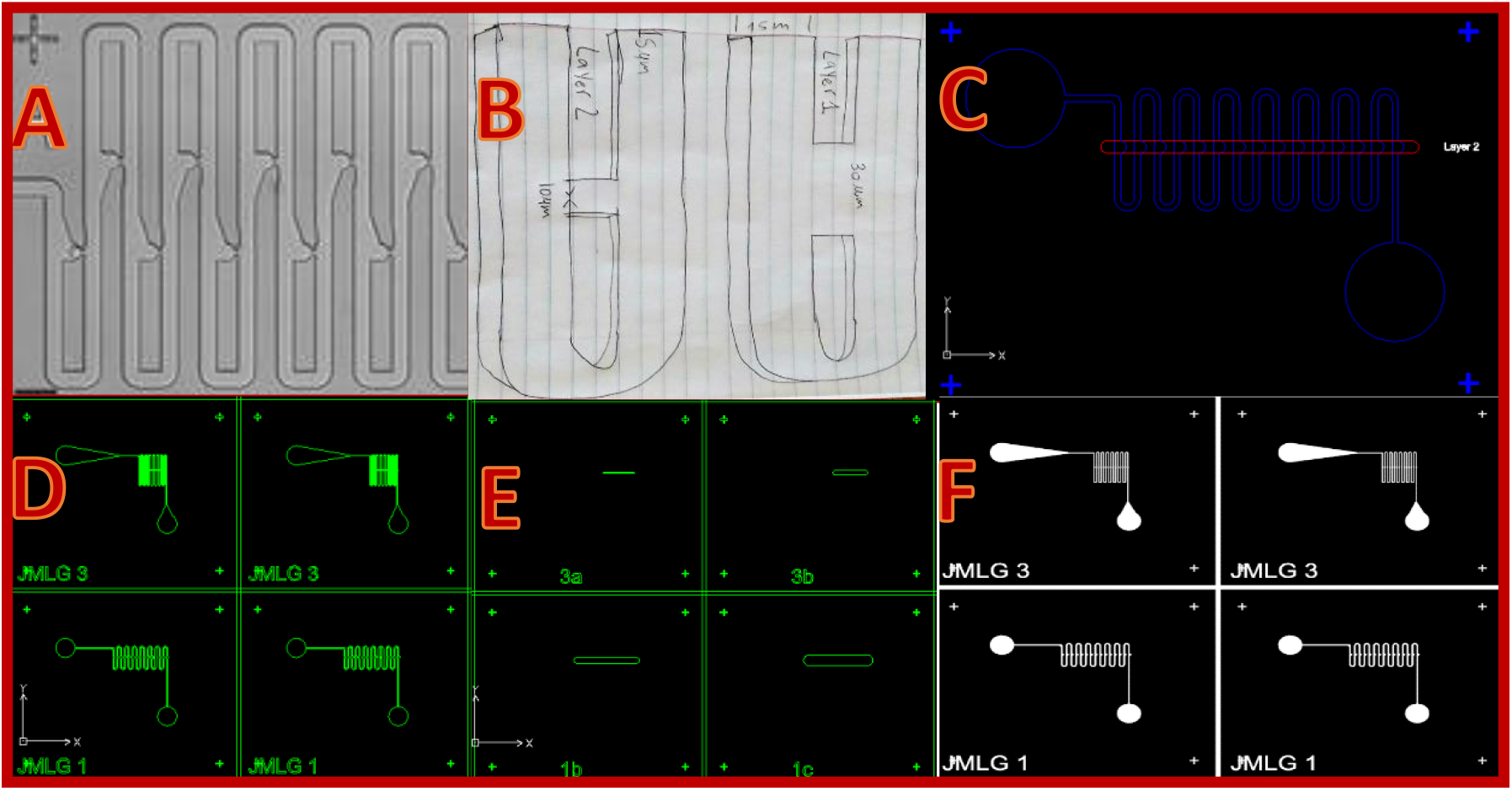
Photomask Design.

#### 2.1.2 Multilayer Photolithography

A 5um thick layer of SU-8 2002 (obtained by dilution of SU-8 2035), was spin coated onto a 4 1/2” diameter pre-cleaned silicon wafer. The wafer was prebaked at 120°C for 5 min to enhance adhesion of the photoresist resin (Figure 2 A), then loaded into the spin coating device. Adding 6 ml of photoresist onto the wafer using a syringe and spun at 200 RPM for 45 seconds (Figure 2 B). The wafer was baked at 95°C for 6 min. Marks for aligning two photomask were added (explained in detail in next section). The wafer and photomask were placed into a UV Kube exposure system and exposed at 23.4 mW/cm^2^ for 11 seconds, giving an exposing dose of 172 mJ/cm^2^ (Figure 2 C). The wafer was baked again at 95°C for 6 min. A 15um thick layer of SU-8 2007 (obtained also by dilution of SU-8 2035), was spin coated, adding 5ml of photoresist resin and spun at 3000 RPM for 45 seconds. The second layer was also prebaked at 95°C for 6 min. The second photomask was aligned to match the first one (explained next section), and exposed to UV light for 11 seconds (172 mJ/cm^2^). The wafer was baked after UV exposition at 95°C for 5 min. The wafer was submerged in SU-8 developer for 2 min, and finally heat-treated in an increasing ratio of 5°C per minute until reach 120°C for 30 min. (Michrochem)(Dirk, 2017)

**Figure 2.**
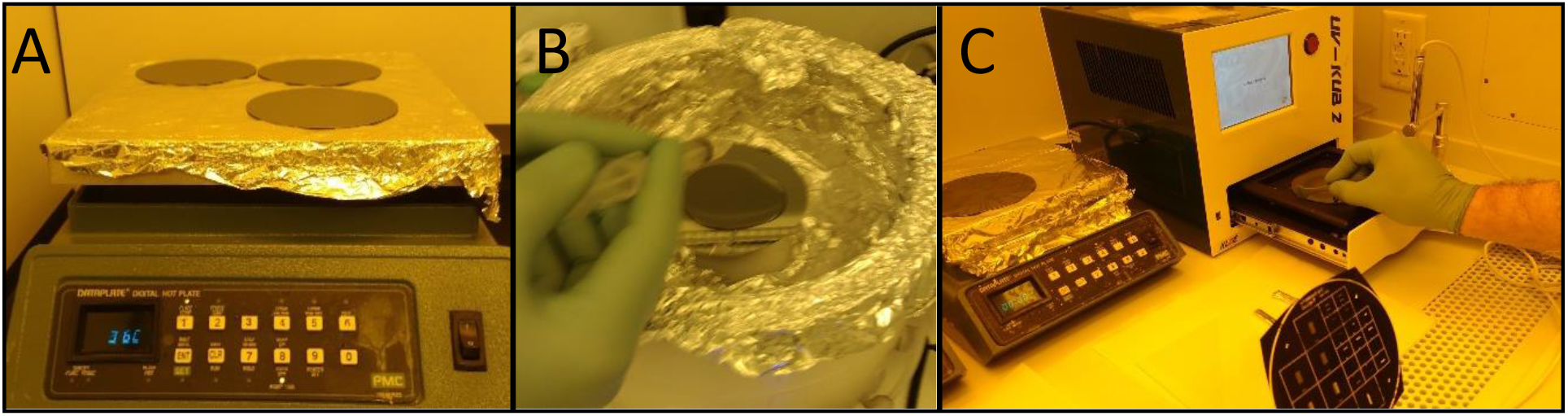
Photolithography Process. Note: Yellow light was used inside clean room to avoid undesired photo-crosslinking of resin.

#### 2.1.3 Aligning Photomasks (Layer 1 and 2 UV crosslinking)

The first step was to put together both photomask and align the layer 1 and layer 2 by eye, using a microscope over a visible mark that forms when both layers are correctly aligned (4 corners) (Fig 3 A). Next procedure was to cut a **“V”** shape form over the four vertex of photomask 1 and 2 clamped together (Fig 3 B). Then small tape pieces were glued to the wafer with photolithographic resin layer 1 on the non-resin area, and marked using a fine tip marker over the aligned photomask 1 before exposing to UV light (Fig 3 C).

**Figure 3.**
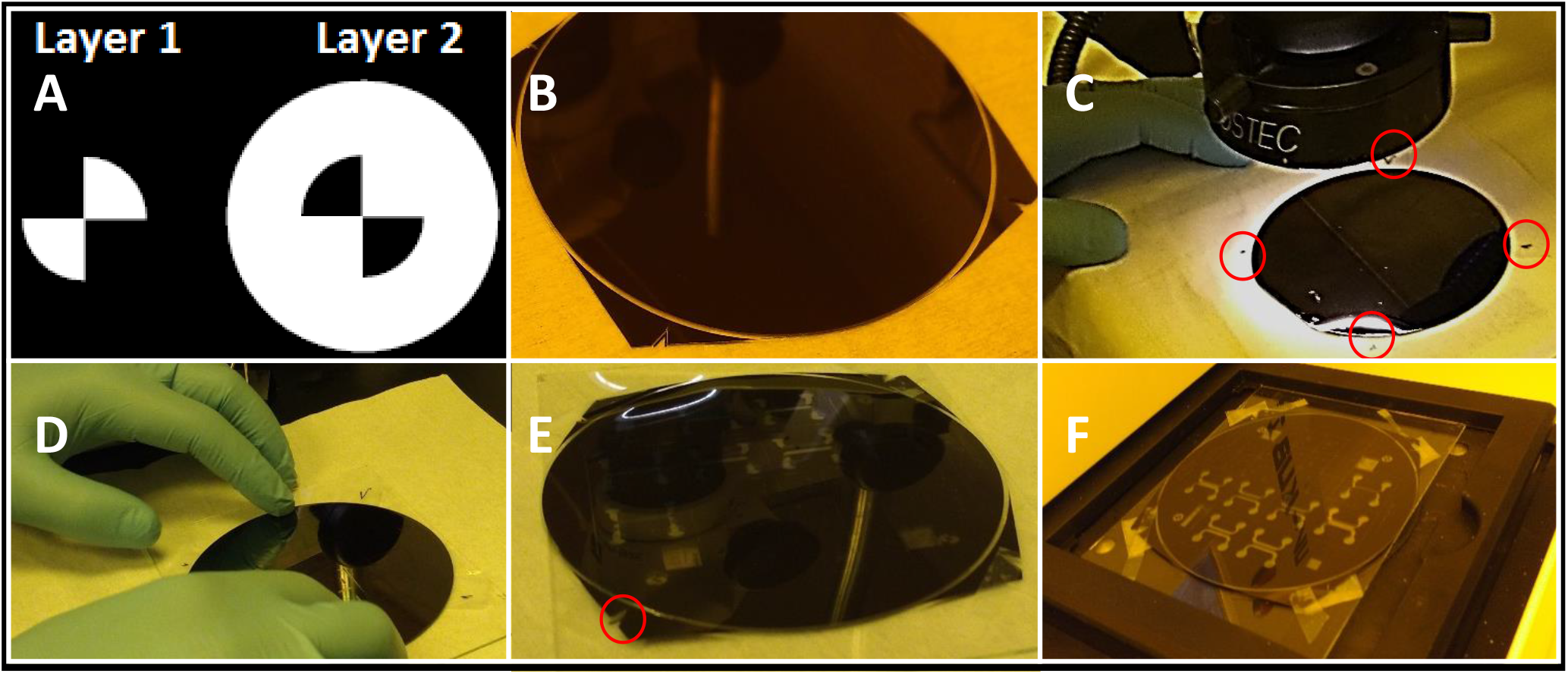
Photomask Aligning Process.

After UV process the marks were covered with tape before pouring the photolithographic resin layer 2 on wafer. When wafer was ready for UV process on layer 2, the cover tapes on marks were removed (Fig 3 D), and photomask 2 was aligned by eye, trying to match the marks on wafer (Fig 3 E). Finally, the aligned photomask 2 was stuck with tape to the wafer and exposed to UV light (Fig 3 F).

#### 2.1.4 Checking results of SU-8 layers on wafer

Once the photolithographic process has finished, was necessary to corroborate the printed layers before continue to soft lithography. Two main checkpoint were critical for proceed to next step: the aligning of the two photomask and thickness of the layers. For this purpose, the **Dektak 3 surface profilometer** was used (Fig 4 A). The profilometer showed layer thickness of 5 and 15 micrometers respectively (Fig 4 B). The alignment of the two layers varies from 50 to 100 micrometers, based on the measures of two repeats of this alignment technique, using as a reference point, structures repeated on both layers (Fig 4 C), and structures existing in different layer but that should overlap (Fig 4 D).

**Figure 4.**
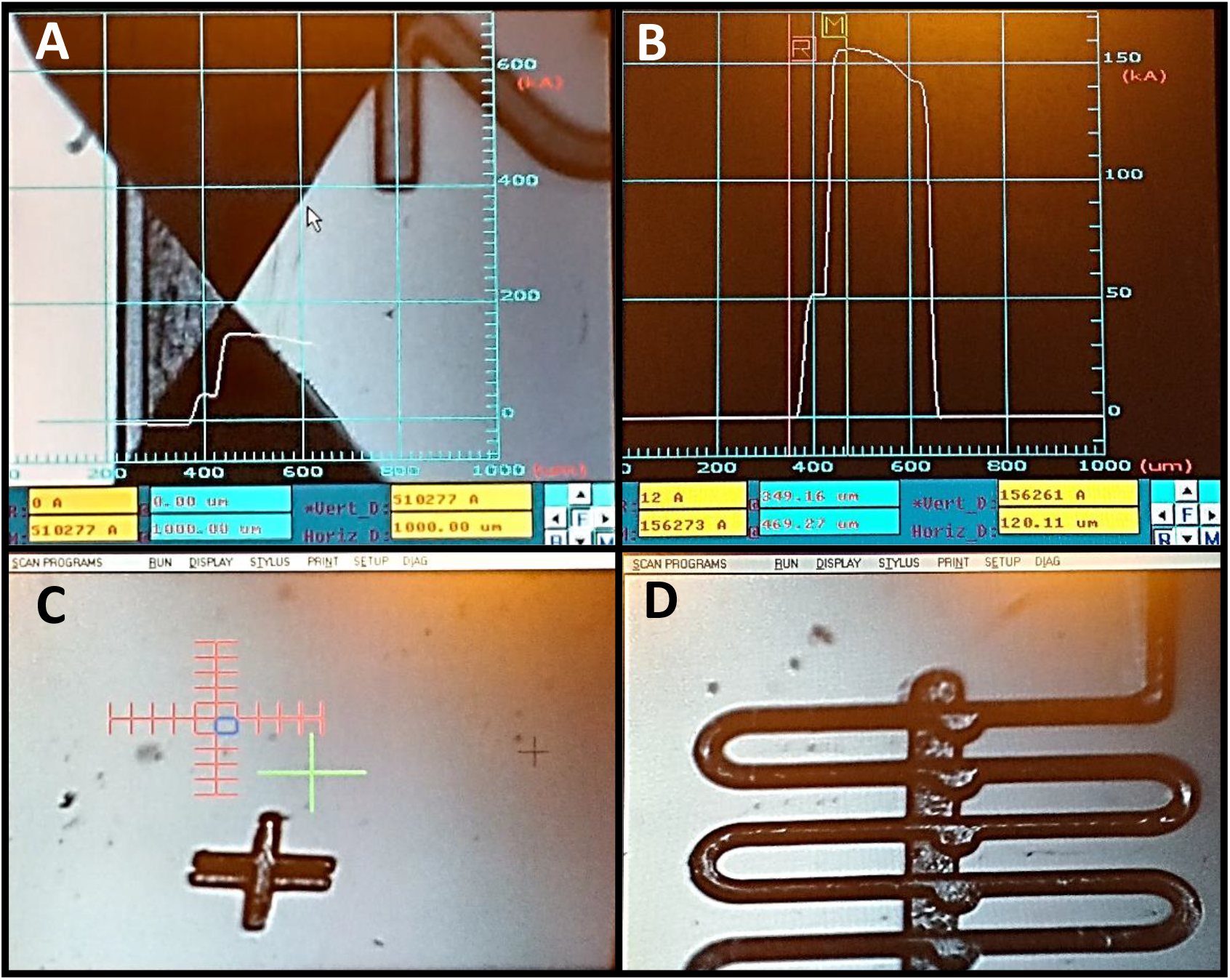
Profilometer: Thickness and alignment analysis.

### 2.2 Soft-Lithography

Soft lithography can be viewed as a complementary extension of photolithography, and is considered part of a family of techniques for fabricating or replicating structures using elastomeric materials most notably using polydimethylsiloxane (PDMS). (Qin et al., 2010) This method has some advantage as the low cost for replication and scaling into mass production of microdevices, and the autosealing property, that can turn into closed chambers after proper treatment of the material.

#### 2.2.1 Casting on PDMS and Plasma Bonding

The wafer with the litography desing on it, was fluorinated by vapor deposition. This procedure facilitates mold release by making hydrophobic the treated surface, preserving the SU-8 pattern for longer time.

Sylgard 184 was used as casting elastomer material (PDMS), which comes as a kit, with Part A (monomer) and Part B (cross linker). It was mixed at a 10:1 ratio, after complete mixing of components, the elastomer was putted into a vacuum chamber for 1 hour for degas the compound and extract all the trapped air bubbles (Fig 5 A). Next procedure was to pour the PDMS into a petri dish containing the silicon wafer; the dish was baked at 60°C overnight. The PDMS was cut of around the wafer and peeled off. The PDMS mold was spliced into individual devices using a razor blade (Fig 5 B). All the inlets ports in devices (∼ 1mm diameter) were punched using a biopsy punch (Fig 5 C).

**Figure 5.**
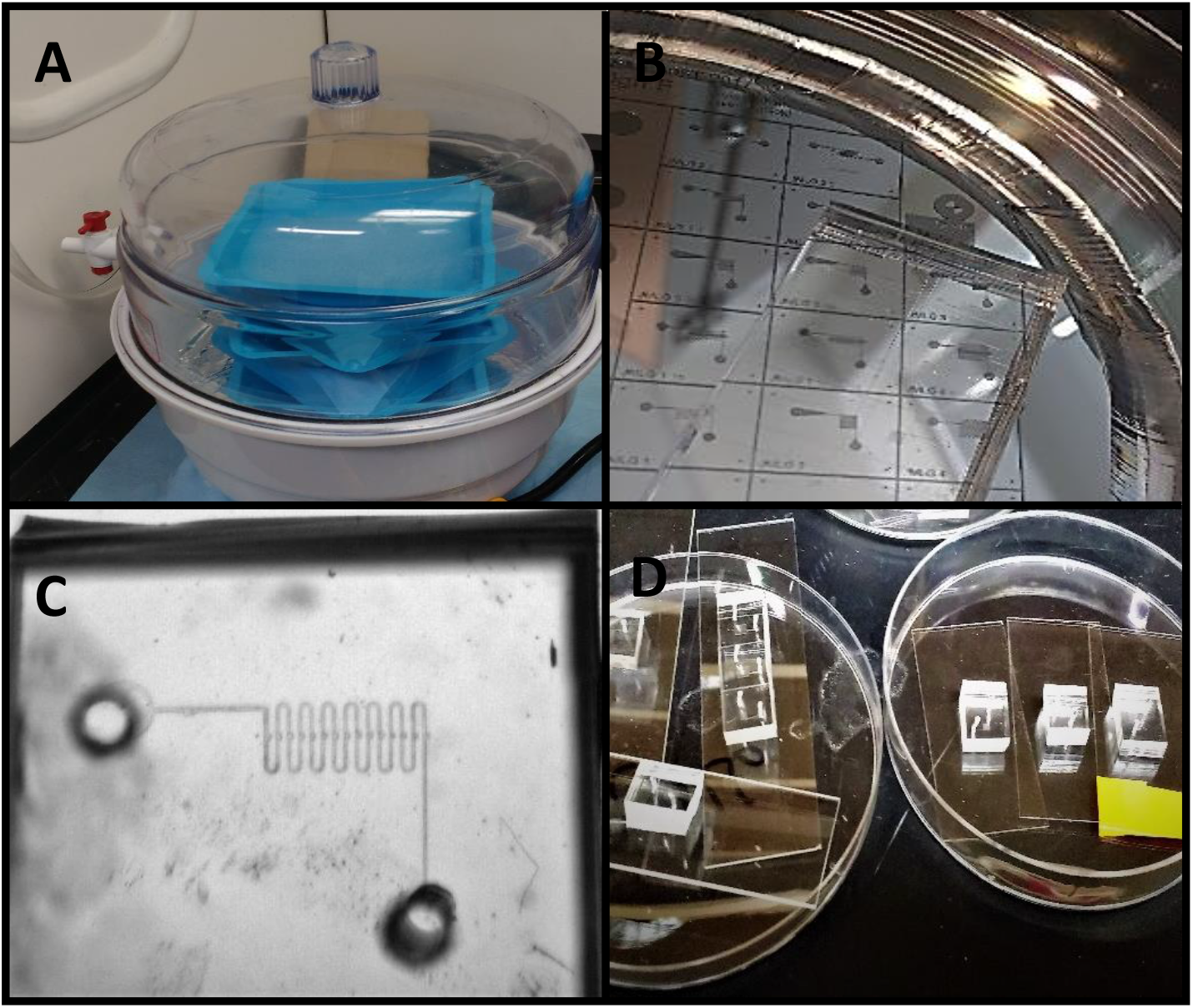
PDMS casting and plasma bonding.

The individual PDMS devices and glass slides were treated with oxygen plasma for 30 seconds. The plasma treated devices were then bound to the treated glass pieces (Plasma Bonding) by inverting the device onto the treated glass surface and applying slight pressure for 10 seconds (Fig 5 D).

#### 2.2.2 Tubing and fluid system

The inlet/outlet of the devices were connected to a tube of electrospun polyethylene terephthalatepolyurethane (PET-PU) from BioSurfaces Inc (Fig 6 A). A 50 cc syringe reservoir suspended at 8” was connected to the inlet port and the outlet port was connected to a receptacle at 1” over the microfluidic device. The plunger of the syringe was removed and the syringe was filled with tap water and yellow food dye, using Gravity Flow System for moving the fluid through the device (Fig 6 B).

**Figure 6.**
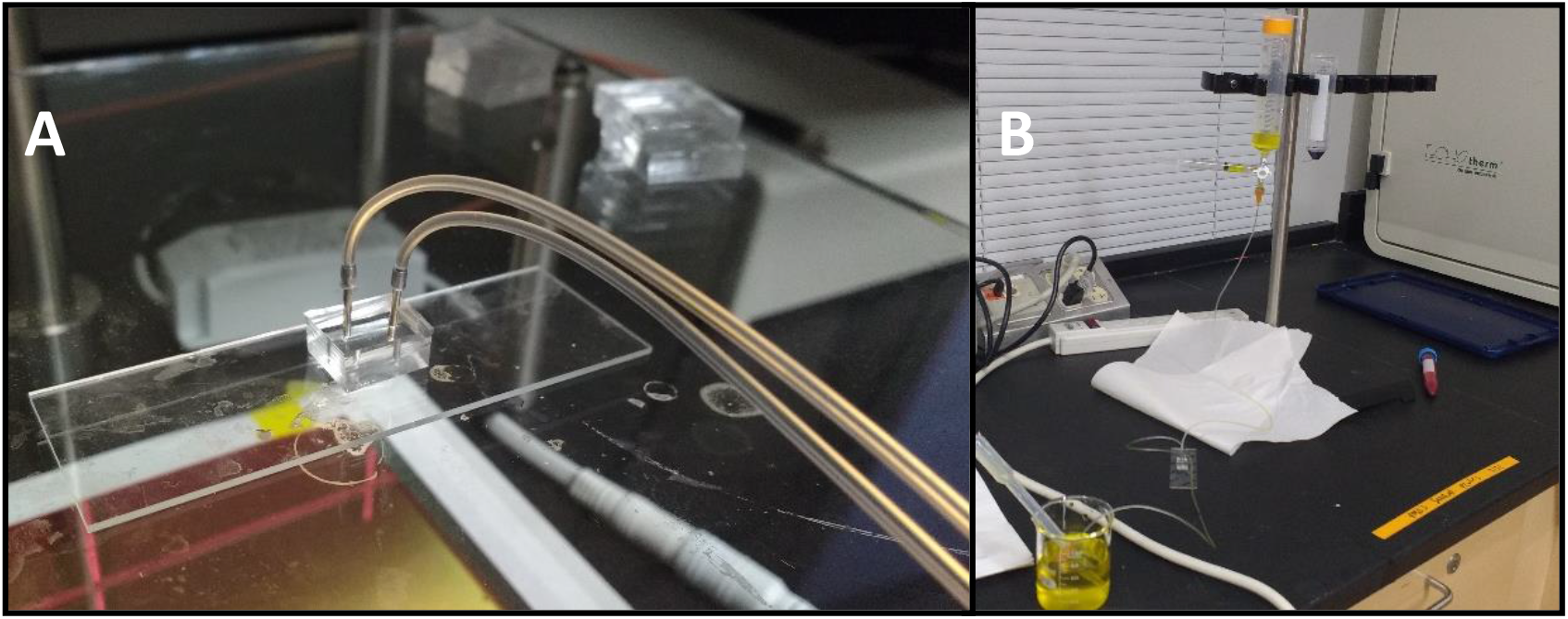
Microfluidic device tubing system.

#### 2.2.3 Flow measuring and cell trapping test

After setting the flow system and corroborating the permeability of the connecting flow through chambers (layer 1 permeability; Fig 8 A), the next procedure was cell trapping test. 5 ml of bovine whole blood (Lampire biomedical) were centrifugated at 2,200 RPM for 10 min, to separate the red blood cells (RBC) from plasma (cell diameter ∼ 5 to 8 um). 1 ml of RBC was incorporated into the flow system connected below the yellow dye (Fig 7 A), and perfused to fill the conecting tube until 50% (Fig 7 B). Then the gravitational flow described in the previous section, was allowed to drive the speed of flow, obtanining the presence of water (yellow) and RBC in the microdevice (Fig 7 C). However, at gravitational speed rate, there was not possible to differentiate individual cells crossing the device or if cells were present in trap chamber. The tubing system connected to the inlet port was readjusted at 4” height and a 45ml reservoir filled with water (also with yellow dye) was connected to the outlet port, slowing down the flow speed allowing cells to enter into microfluidic device at lower speed (Fig 7 D), and being captured into the chambers (Fig 8 B).

**Figure 7.**
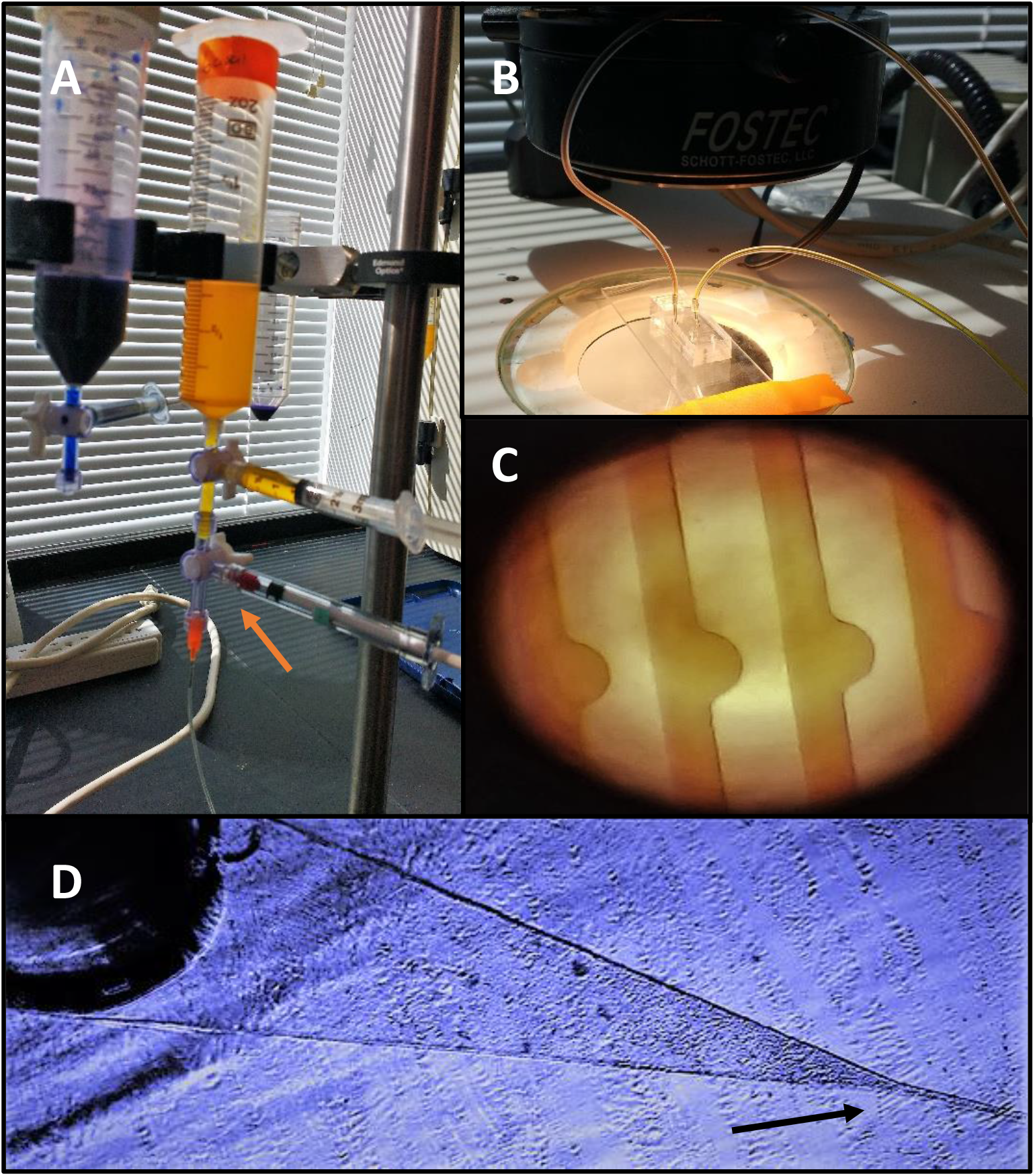
Cell trapping test.

**Figure 8.**
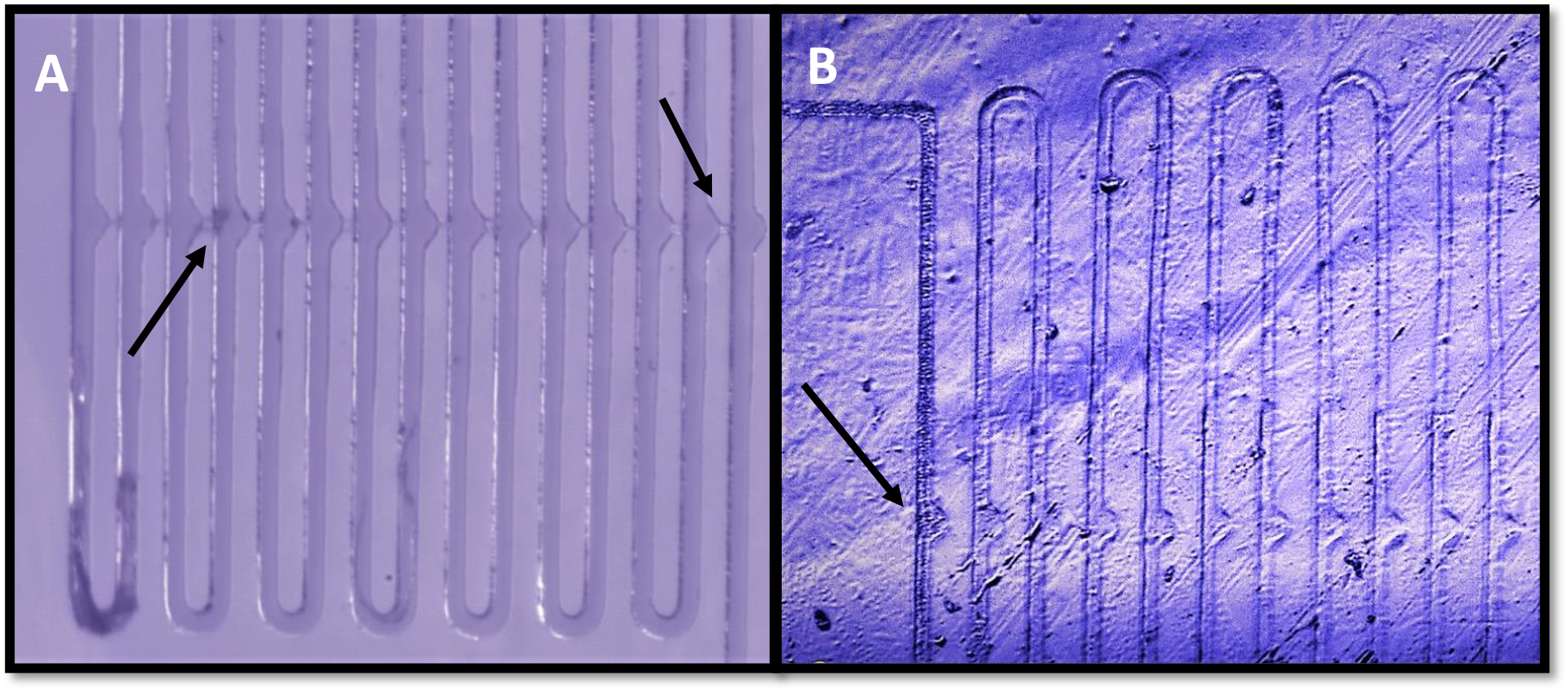
Cell trapping chambers.

## 3. Results

The overall goal of this study was the fabrication of a hydrodynamic cell trap device, using low-cost methods that could replicate the results obtained by kimmerling team. This device utilizes an array of hydrodynamic traps chambers within a fluidic design optimized to capture and culture cells for multiple generation’s (Fig. 8 B). However, from the nine variations on design, only four were able to work as intended or trap cells (Fig. 9).

**Figure 9.**
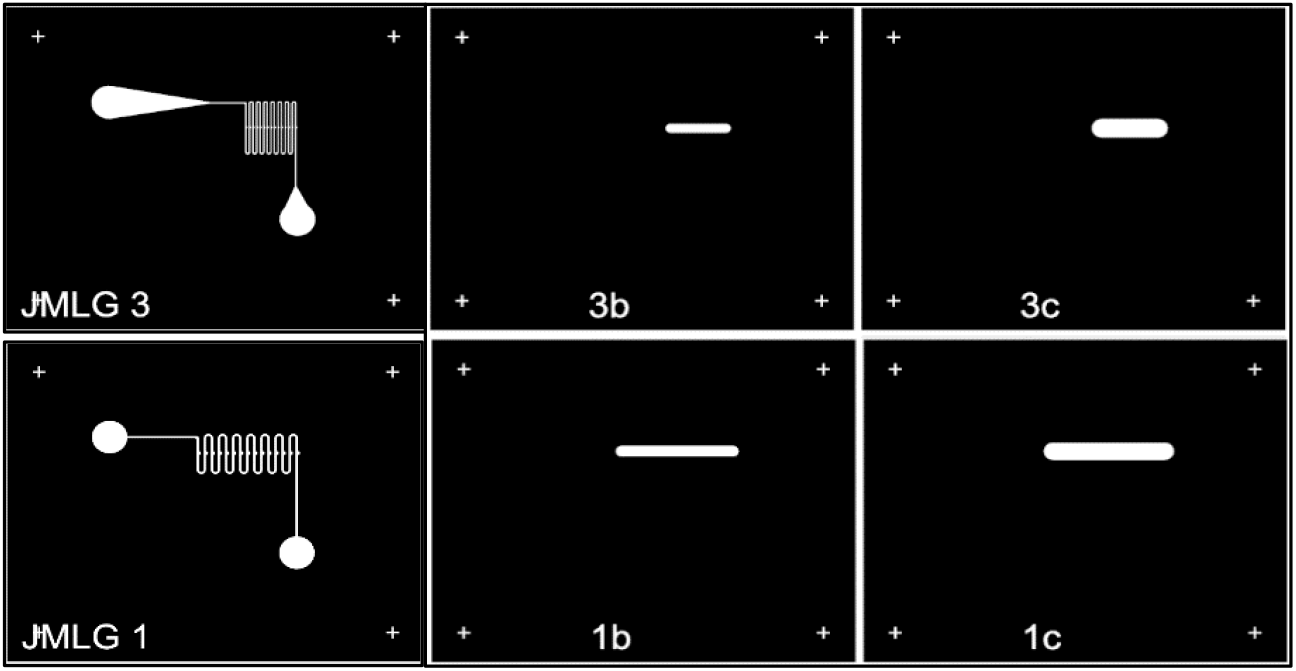
Working Microfluidic devices.

After determine the functional devices, the permeability tests began, looking for the movement of fluids between chambers and layers. A 5 um height channel connected each trapping chamber (15um height). However, it was difficult to observe the connecting layer at microscope, for confirming their presence, a two dye experiment was conducted. Understanding that depending on velocity of the fluid crossing the device, this will access the 5 um channels (more resistance) or continue over the main channels (50 um width and 15 um height). First, a yellow dye by gravity flow at 4” was introduced into system, after 5 minutes a new dark purple dye was introduced changing velocity by gravity flow at 8”; a picture was taken at that moment (Fig 10), allowing to observe the boundaries of the 5um channels (squared in orange). For testing, the permeability on both sides of the device, the water flux was inverted at same speed that introducing yellow dye with Deionized water (DI water, green arrow), showing a completely clearance of dye of the device.

**Figure 10.**
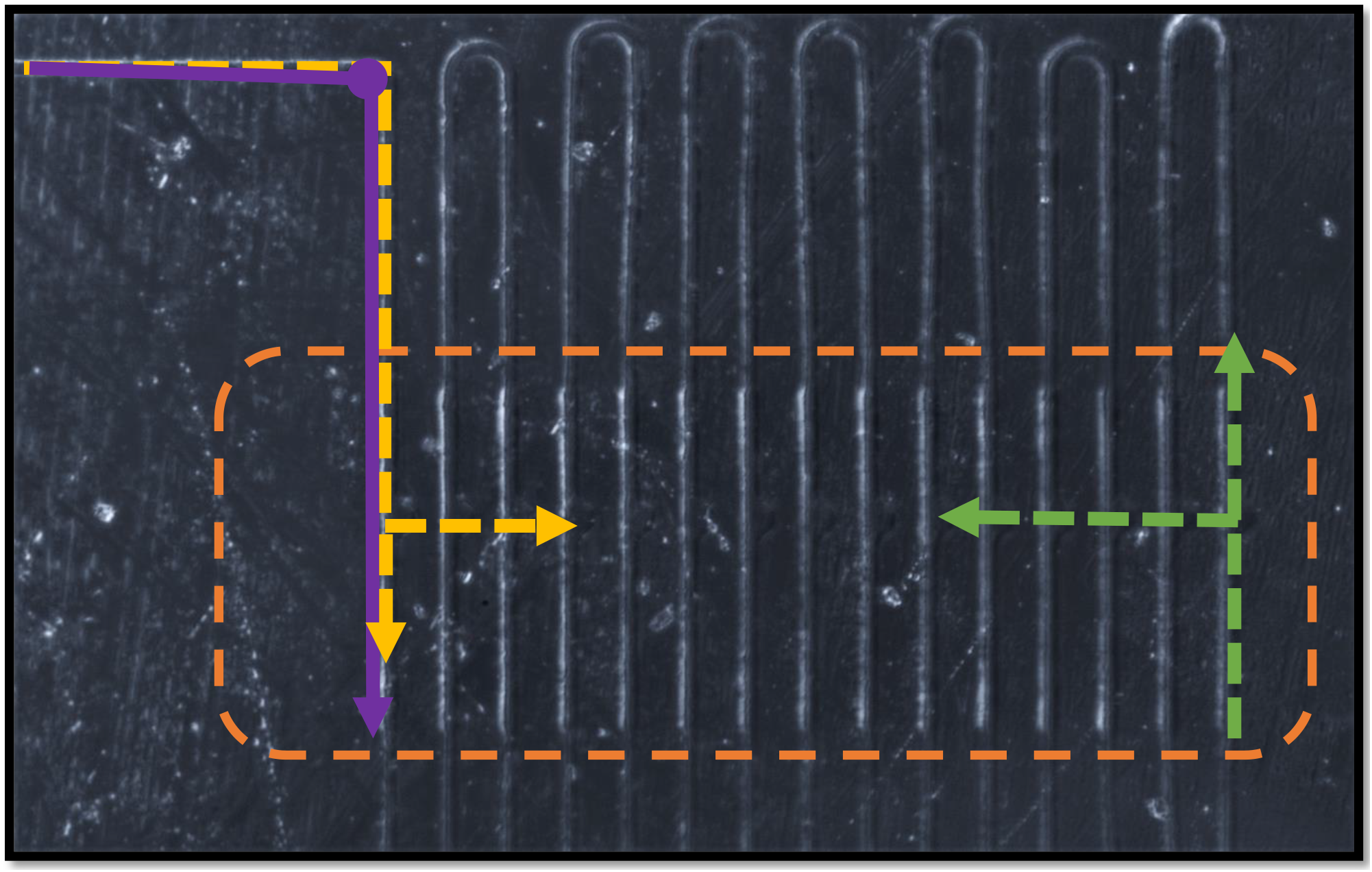
Hydrodinamic behavior of fluids (2 layers)

After assuring the correct functioning of device, cells were seeded, but in first tests, cells move too quickly to be observable (Fig 11 C), https://youtu.be/F1XZaYxAS2s. For that reason to determinate the actual flow speed that was entering into device to be adjusted as needed, the Haze-Williams equations were used. Three speed flows were tested, corresponding to different height of the water reservoir (8”,6”,4”). The experimental confirmation looking at the microscope of the behavior of the red cells determined that the optimal speed was 0.189 m/s corresponding to the height of 4” (Fig 11 A), also showing that using the same speed on reverting flux will be optimal https://youtu.be/hOyIc9fvdBw.

**Figure 11.**
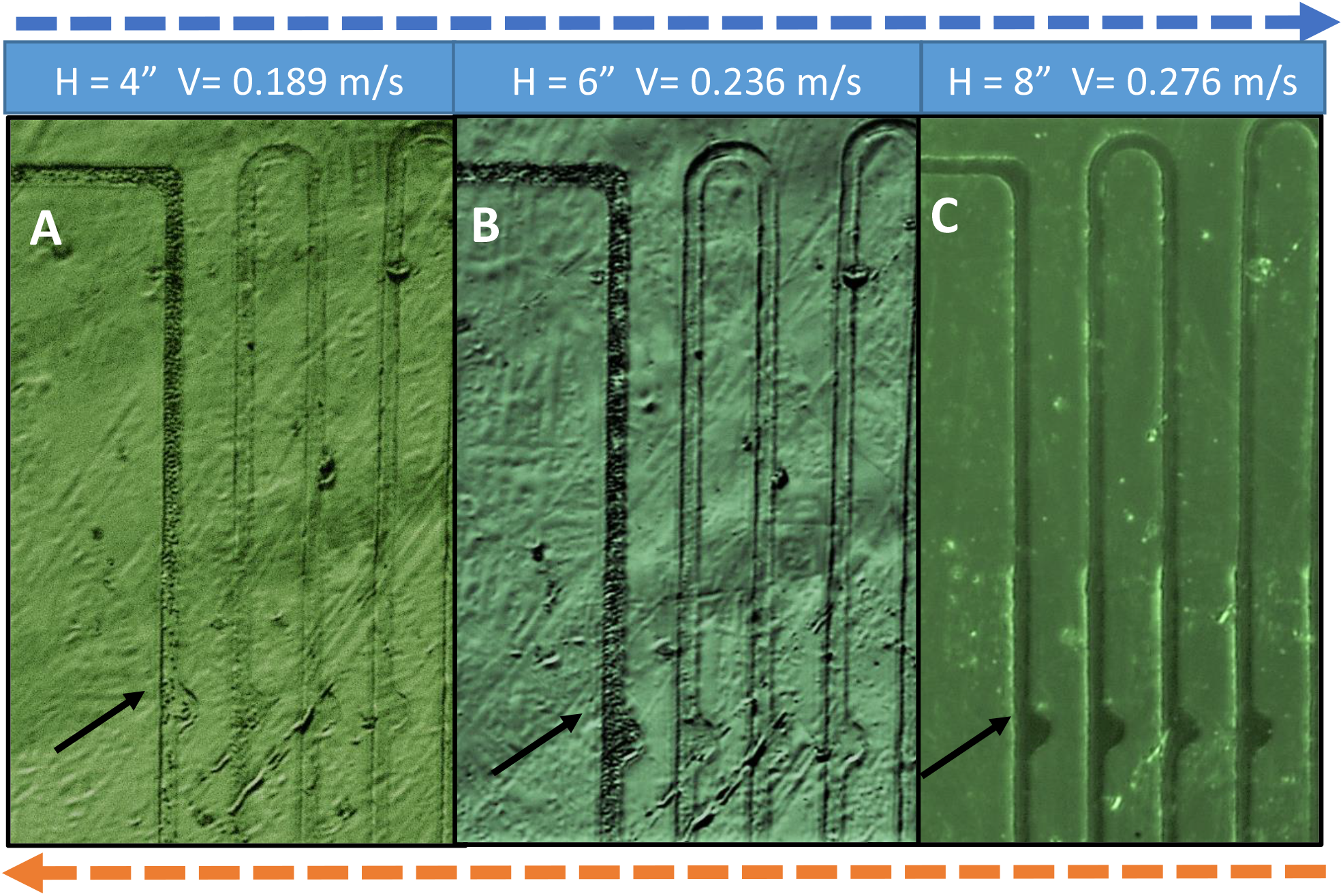
Variation of flow speed for cell seeding.

### Hazen-Williams equations

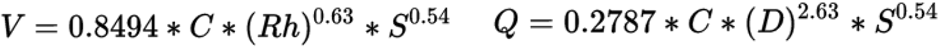

**Di** = Inner diameter = **0.001 m Rh** = Hydraulic Radius = Di / 4 = **0.00025** m

**V** = Velocity of water inside tube [m/s]. = **0.275 m/s at 8”; 0.236 m/s at 6”; 0.189 m/s at 4”**

**H** = H1 =**8” (0.2m)**. H2 =**6” (0.15m)**. H3 =**4” (0.1m)**.

**S** = Slope ((Height of container inlet – Height of container outlet)/length of tubing) [m/m]

**D** = Distance = **0.95 m**.

**C** = Polyethylene coefficient = **140**.

Once the correct speed was established, the analysis of chambers started, observing that based on the design chambers, they tend to trap more than one cell at time, by multiple repetition of the loading and cleaning process (DI water), around 5 to 10 RBC cells were trapped at every chamber by cycle (Fig. 11 A). After inverting fluid direction, it was possible to release trapped cells, and reoriented as necessary, if multiple passes of inverting and normalizing flux cells tends to form well-organized lines.

## 4. Discussion

The fabrication of a microfluidic device involves many steps and requires a minimum equipment that could be expensive. The complexity of the devices has a great impact in the cost and effort in production. However, taking advantage of new ways to approach the fabrication process, like using a multilayer technique, economical limitations can be overpassed.

This work show that the photolithographic process is not complicated by itself, but obtaining the accuracy on what is planned it is the complicated part. For the thin layer (5 um) the use of an SU-8 2002 resin obtained by dilution of SU-8 2035, could lead to height inaccuracy, because the aimed height was 3 um. Something similar happened at the second layer height were SU-8 2007 was obtained also by dilution method, in this case obtained layer height was 15 um and originally intend was 7.5 um. The dilution method was selected based on prices of the other two resins (2002 and 2007), that are exact dilutions of SU-8 2035. Despite of the difference on expected against obtained heights the relation between layer 1 and 2 was preserved enough to continue with fabrication of the device.

Another complication arise in the layer aligning process, were extreme accuracy was needed. For that reason only 4 of the 9 designs on photomask worked. Only those devices where an overlapping tolerance were superior to 50um tend to work. However, after multiple attempts at alignment the results could not reach alignment superior to that threshold. With this knowledge on mind, taking on consideration a 50 um inaccuracy on alignment at early stages of photomask design, it is possible to fabricate a multilayer pattern.

When the photolithography process was completed, the soft-lithography did not present technical challenges, however, observing the correct proportions of the components for preparing the PDMS, and baking times is imperative to correct stamping of the pattern on the soft gel. For plasma bonding process, is necessary to clean carefully the surfaces of materials to avoid leaking on the resulting device.

For testing the cell trapping chambers, once the optimal fluid speed for the device was calculated, the selection of red blood cells (RBC) was not arbitrary; it was based on their easy procurement and no need for culturing over multiple tests, also they are easy to observe without special live staining. Based on the size of RBC (5 to 7 um) the number of cells trapped is consistent around 5 to 10, however seeding bigger cells with special staining this design could trap a fewer number of cells in each chamber and will be easier to see them.

## 5. Conclusions

It is possible to fabricate a functional two-layer microfluidic device capable of trap and posteriorly release cells using low-cost resources, taking advantage of simple techniques for alignment and imaginative design for overcome limitations on miniaturization.

These devices could be escalade up for multiple applications, varying the cells seeded, and conditions of functioning, for purposes as different as cancer mitotic studies or understanding the bacterial growth in small colonies.

## References

Alvaro, M., Aaron, J. F., & Shuvo, R. (2006). Fabrication of multi-layer SU-8 microstructures. Journal of Micromechanics and Microengineering, 16(2), 276.

Choi, S., & Park, J.-K. (2010). Two-step photolithography to fabricate multilevel microchannels. Biomicrofluidics, 4(4), 046503. doi:10.1063/1.3517230

Dirk, A. (2017). Photolithography using SU8 Photoresist STANDARD OPERATING PROCEDURE. WPI protocol. Biomedical Engineering. Worcester Polytechnic Institute.

Jaeger, R. C. (2002). Introduction to microelectronic fabrication (2nd ed. Vol. 5.;5;). Upper Saddle River, N.J: Prentice Hall.

Kimmerling, R. J., Lee Szeto, G., Li, J. W., Genshaft, A. S., Kazer, S. W., Payer, K. R., … Manalis, S. R. (2016). A microfluidic platform enabling single-cell RNA-seq of multigenerational lineages. Nature Communications, 7, 10220. doi:10.1038/ncomms10220 http://www.nature.com/articles/ncomms10220#supplementary-information

MicroChem, SU-8 2000 Permanent Epoxy Negative Photoresist PROCESSING GUIDELINES FOR: SU-8 2025, SU-8 2035, SU-8 2050 and SU-8 2075,” ed, pp. 1-5. http://www.microchem.com/pdf/SU-82000DataSheet2000_5thru2015Ver4.pdf

Lin, L., Chu, Y.-S., Thiery, J. P., Lim, C. T., & Rodriguez, I. (2013). Microfluidic cell trap array for controlled positioning of single cells on adhesive micropatterns. Lab on a Chip, 13(4), 714–721. doi:10.1039/C2LC41070B

Qin, D., Xia, Y., & Whitesides, G. M. (2010). Soft lithography for micro- and nanoscale patterning. Nat. Protocols, 5(3), 491–502.

